# Glycan-specific IgM is critical for human immunity to *Staphylococcus aureus*

**DOI:** 10.1101/2023.07.14.548956

**Authors:** Astrid Hendriks, Priscilla F. Kerkman, Meri R.J. Varkila, Jelle L.G. Haitsma-Mulier, Sara Ali, Thijs ten Doesschate, Thomas W. van der Vaart, Carla J.C. de Haas, Piet C. Aerts, Olaf L. Cremer, Marc J.M. Bonten, Victor Nizet, George Y. Liu, Jeroen D.C. Codée, Suzan H.M. Rooijakkers, Jos A.G. van Strijp, Nina M. van Sorge

## Abstract

*Staphylococcus aureus* is a major human pathogen but the immune factors that protect against it remain elusive. In particular, high opsonic IgG titers achieved in preclinical *S. aureus* animal immunization studies have consistently failed to translate to protection in human clinical trials. Here, we investigated the antibody responses to a conserved surface glycan, Wall Teichoic Acid (WTA). IgM and IgG antibodies specific to WTA were universally present in plasma from healthy individuals. Functionally, WTA-specific IgM outperformed IgG in opsonophagocytic killing of *S. aureus* and conferred passive protection against *S. aureus* infection *in vivo*. In the clinical setting, WTA-specific IgM responses, but not IgG responses, were significantly lower in *S. aureus* bacteremia patients compared to healthy individuals, correlated with mortality risk and showed impaired bacterial opsonization. Our findings can guide risk stratification of hospitalized patients and inform future design of antibody-based therapies and vaccines against serious *S. aureus* infection.

## Introduction

The bacterial pathogen *Staphylococcus aureus* is among the leading causes of both community- and hospital-acquired infections in human medicine. Despite significant advances in intensive care management, high rates of morbidity and mortality persist for patients with *S. aureus* bacteremia (1). This problem is further compounded by the emergence of antibiotic-resistant strains, particularly methicillin-resistant *S. aureus* (MRSA), which hinders both treatment and prophylaxis efforts (2). To improve future clinical outcomes, improved risk stratification as well as the development of alternative strategies, such as vaccines and immune-based approaches, is imperative. However, a precise understanding of the immune correlates responsible for protection against *S. aureus* remain elusive.

Neutrophils play a critical role in eradicating *S. aureus*, as demonstrated by human genetic studies and animal experiments showing that defects in neutrophil responses predispose individuals to infection (3, 4). To effectively combat *S. aureus*, neutrophils rely on additional immune factors, such as opsonic antibodies and complement. Ig-mediated complement activation leads to the deposition of C3b, enhancing bacterial uptake and killing by neutrophils and promoting neutrophil recruitment through the release of potent chemotactic factors such as C5a (5, 6). Despite the success of opsonic IgG antibodies against other pathogens, such as pneumococci, meningococci, and *Haemophilus influenzae* b (Hib), efforts to identify phagocytosis-enhancing IgG targeting conserved surface structures for *S. aureus* have faced challenges (7–9). Therapeutic antibodies and vaccines targeting surface-exposed antigens that showed promise in preclinical animal models have consistently failed in human clinical trials, highlighting the need for antibody profiling studies in relevant patient cohorts to improve predictive value (7, 8).

One of the most prominent *S. aureus* virulence factors is staphylococcal protein A (SpA), an abundant and universally-expressed surface molecule. SpA effectively inhibits Fc-mediated IgG effector functions, including complement activation and Ig-mediated phagocytosis (10, 11), which may explain the limited effectiveness of IgG in real-world studies. However, IgG is not the only antibody isotype capable of boosting neutrophil activity. IgM, the third most abundant isotype in blood, exhibits complement activation capacity (12). Importantly, SpA does not bind the Fc region of IgM, suggesting that its opsonic capacity remains unaffected by SpA. Furthermore, IgM shows enhanced binding to structures with repetitive epitopes, such as microbial carbohydrates, due to its higher avidity compared to other Ig isotypes (13). Despite these potential benefits, the role of *S. aureus*-specific IgM in host defense has received limited research attention.

*S. aureus* wall teichoic acids (WTAs) are conserved cell-wall anchored glycopolymers and a dominant target for opsonic antibodies, with as much as 70% of the total surface-directed IgG pool binding to WTA (14). However, *S. aureus* WTAs exhibit distinct structural variation due to glycosylation with *N*-acetylglucosamine (GlcNAc), impacting immune interactions and antibody recognition (15, 16). Three WTA glycotypes have been identified, each requiring a dedicated glycosyltransferase enzyme (TarM, TarS or TarP) to add the GlcNAc to the ribitol phosphate WTA backbone subunits in an α1,4-(TarM), β1,4-(TarS) or β1,3-(TarP) position (17–19). All *S. aureus* isolates possess the *tarS* gene, while approximately 40% of strains co-express *tarM* or *tarP* (20). The specific WTA glycoprofile of a particular *S. aureus* strain is determined by the presence of corresponding *tar* genes, enzymatic activity, and environmental conditions, leading to glycovariation (21). The diversity in S*. aureus* WTAs holds important implications for immune responses and potential antibody-based therapeutics.

IgG antibodies, predominantly IgG2, have been detected against all three WTA-GlcNAc glycotypes in serum from healthy donors, with a higher reactivity toward β-1,4-GlcNAc and β-1,3-GlcNAc compared to α-1,4-GlcNAc-WTA (15). These antibodies have demonstrated enhanced complement deposition and neutrophil killing of SpA-deficient *S. aureus* (22–24). Despite the widespread reactivity of antibodies to WTA in healthy individuals, their potential contribution to protection against severe *S. aureus* infections in humans remains unexplored. In this study, we used synthetic WTA-oligomers that mimic *S. aureus* WTA to investigate human systemic antibody responses to the three main *S. aureus* WTA glycotypes. Unexpectedly, our studies revealed a critical role for WTA-specific IgM in immune protection.

## Results

### The antibody repertoire to *S. aureus* WTA in healthy individuals

We employed a bead-based assay, adapted for multiplexing, using synthetic WTA oligomers to study the antibody repertoire (IgG1, IgG2, IgG3, IgM, IgA) targeting specific *S. aureus* WTA glycotypes (Supplementary Figure 1) (15). Analysis of plasma samples from 31 healthy donors revealed specific IgM responses to all three WTA glycotypes in 30 out of 31 (97%) donors (Figure 1a). Furthermore, WTA-specific IgG2 antibodies were detected against at least one WTA glycotype in all donors (Figure 1b). Conversely, only low levels of WTA-specific IgA, IgG1 and IgG3 were observed in approximately half of the donors (Supplementary Figure 2). IgG2 responses to TarS-modified WTA were consistently high and similar to those against TarP-modified WTA (Figure 1b), likely due to cross-reactive antibody clones recognizing β-GlcNAc-WTA (Figure 1c) (15). In contrast, IgM antibodies demonstrated a significant positive correlation in binding to all three WTA glycotypes, indicating the presence of poly-reactive IgM clones that bind GlcNAcylated WTA irrespective of GlcNAc configuration or linkage position (Figure 1d). These observations highlight the extensive and diverse antibody repertoire targeting *S. aureus* WTA in healthy individuals, with potential variations in binding specificities and characteristics across different antibody isotypes (IgG and IgM).

**Figure 1.**
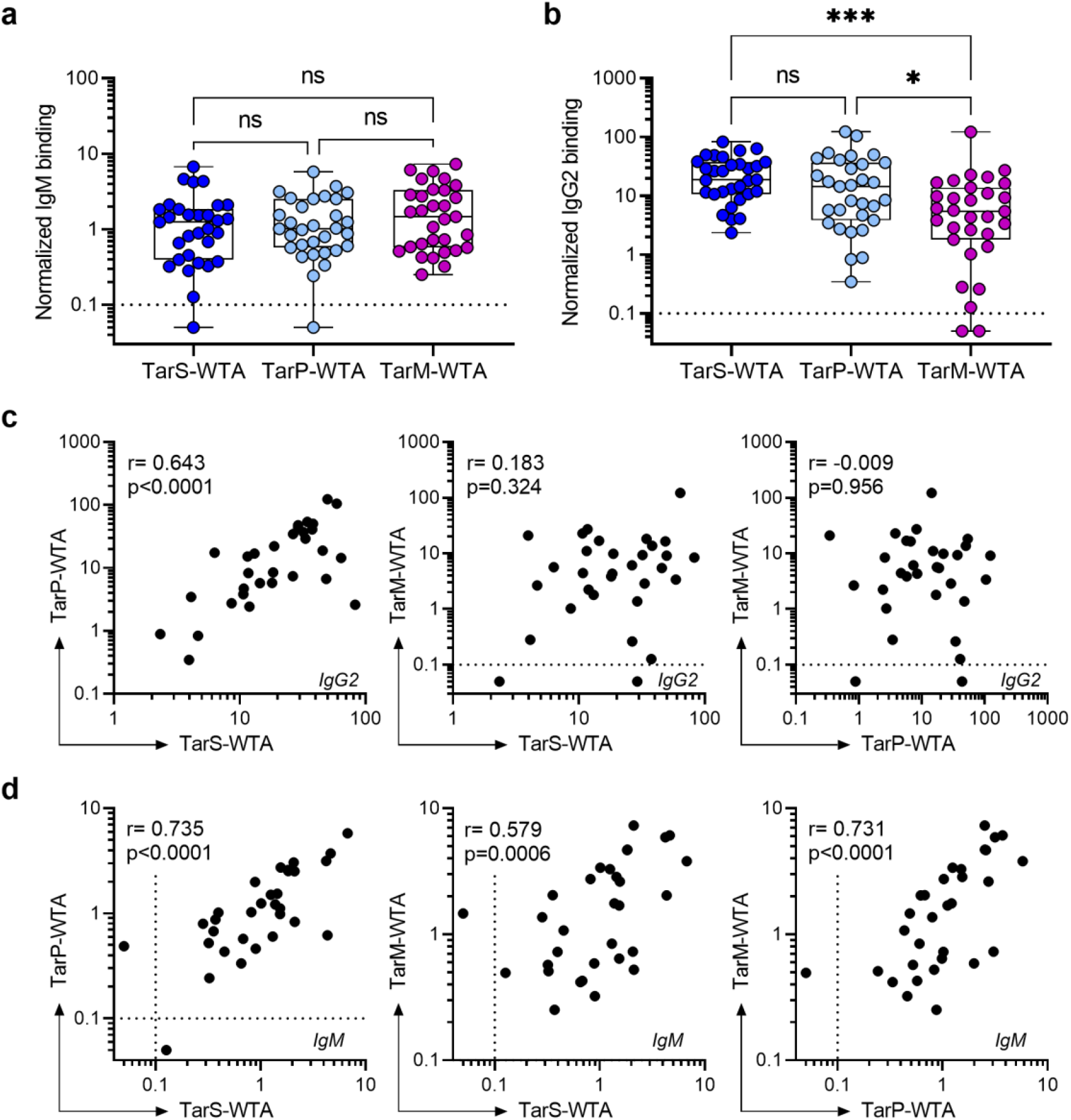
IgM and IgG2 antibody responses against three *S. aureus* WTA glycotypes in healthy subjects. a, b) Normalized binding of (a) IgM and (b) IgG2 to beads coated with TarS-WTA, TarP-WTA and TarM-WTA in plasma from healthy donors (n=31). Boxplots extend from the 25th to 75th percentiles and the line represents the median. c, d) Correlation between binding of (c) IgG2 and (d) IgM reactivity toward distinct WTA glycotypes within individual donors. Each dot represents an individual donor (n=31). Dotted line represents the lower limit of detection, symbols shown below line present extrapolated values. *p < 0.05, ***p < 0.001.

### WTA-specific IgM contributes to protection against *S. aureus* infection *in vitro* and *in vivo*

We postulated that WTA-specific IgM could play a more prominent role in protective immunity against *S. aureus* than IgG2, since the effector functions of opsonic IgG antibodies can be subverted by SpA. To this end, we compared the efficacy of IgM and IgG2 monoclonal antibodies derived from an original IgG1 clone targeting β-GlcNAc WTA (clone 4497) (14, 25). Our results showed that the IgM variant, 4497-IgM, was over 100-fold more effective at inducing complement C3b deposition on *S. aureus* compared to 4497-IgG2 at equimolar concentrations (Figure 2a). Importantly, the presence of surface bound (Figure 2a) or secreted SpA (Figure 2b) did not interfere with IgM-mediated C3b deposition, unlike IgG2. Furthermore, complement deposition by 4497-IgM triggered neutrophil-mediated killing of *S. aureus*, whereas 4497-IgG2 showed no such effect at equimolar concentrations (figure 2c). To evaluate the protective potential *in vivo*, mice were passively immunized with 4497-IgM prior to systemic *S. aureus* infection. The 4497-IgM-treated mice showed significantly lower CFU counts in spleen compared to control groups treated with PBS or a non-specific IgM antibody (figure 2d). These findings strongly support the notion that WTA-specific IgM plays a critical role in conferring protection against systemic *S. aureus* infection, likely by enhancing opsonophagocytic killing of the pathogen by neutrophils, the key effector cells in innate host antibacterial defense.

**Figure 2.**
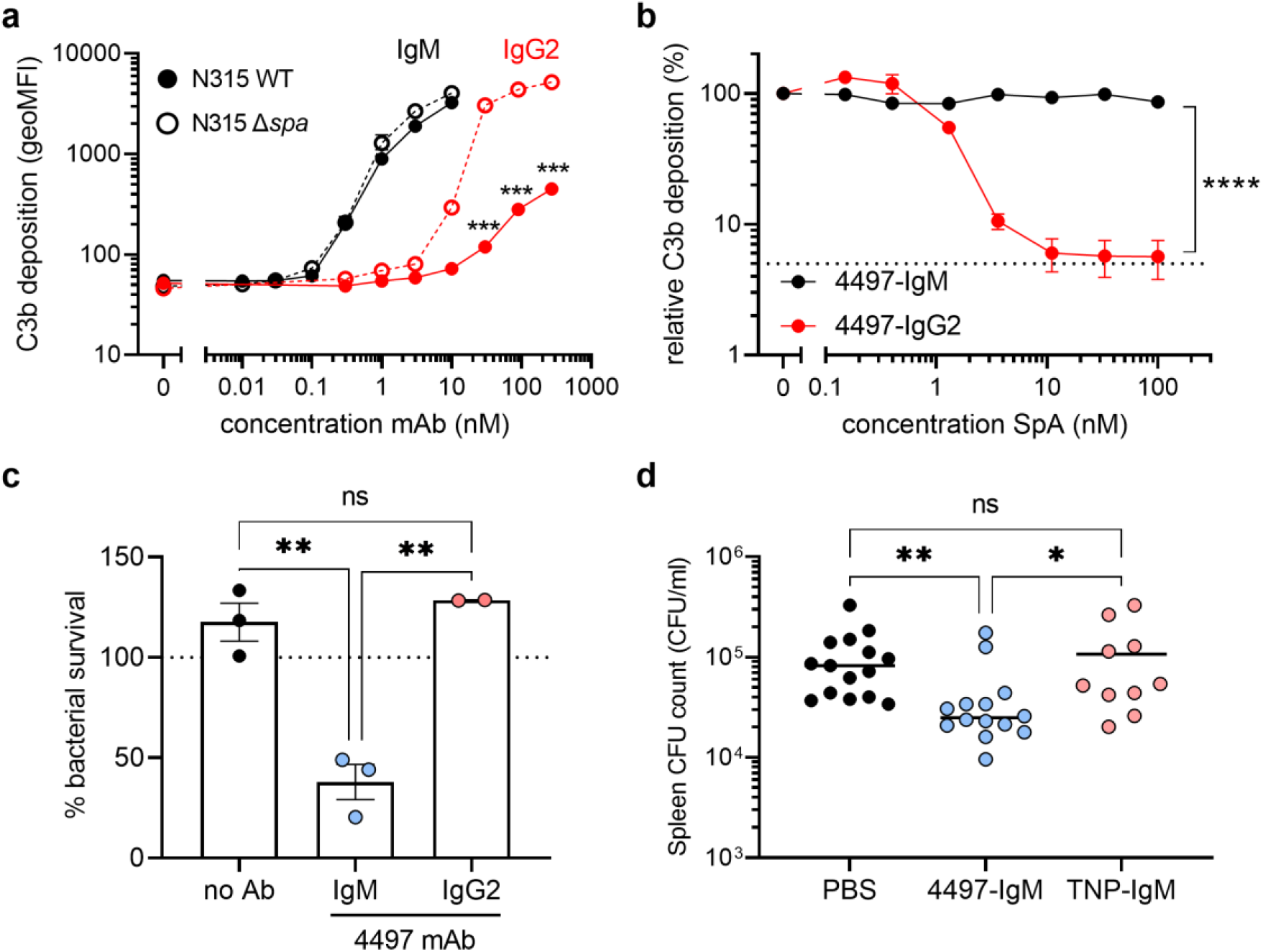
WTA-specific IgM supports immune-mediated clearance of *S. aureus* killing *in vitro* and *in vivo* mouse infection experiments. a) Complement C3b deposition on *S. aureus* N315 wildtype (WT) or staphylococcal protein A-deficient (Δspa) bacteria, pre-opsonized with 4497-IgG2 (0.3-270 nM) or 4497-IgM (0.01-10 nM). b) Relative complement C3b deposition on SpA-deficient *S. aureus* (Newman Δ*spa*/*sbi*), pre-opsonized with 4497-IgG2 (10 nM) or 4497-IgM (1 nM) in presence of recombinant protein A (SpA, 0.15-100 nM). C3b deposition in the absence of SpA was set at 100%, the dotted line represents background C3b deposition without antibody opsonization. c) Neutrophil opsonophagocytic killing (OPK) of *S. aureus* N315 WT, in presence of 10 nM 4497-IgM, 10 nM 4497-IgG2, or no antibody control. Bacterial survival has been normalized over inoculum (dotted line at 100%), data is shown from three independent donors (two donors in case of IgG2) d) Spleen CFU counts from mice (n=10-15) at 24 h post infection with *S. aureus* N315 WT, passively immunized with 30 µg 4497-IgM, anti-TNP-IgM or PBS control. ns= non-significant, *p < 0.05, **p <0.01, ***p < 0.001, ****p < 0.0001.

### Patients with *S. aureus* bacteremia have low levels of WTA-IgM

I*n vivo* animal models have a poor track record for predicting translational success in human vaccine and therapeutic antibody development (26, 27). To avoid overreliance on experimental models, we therefore further investigated the role of WTA-specific IgM antibodies in protective immunity in a cohort of 36 ICU patients with *S. aureus* bacteremia (Table 1) by assessing the natural magnitude of WTA-specific IgM and IgG responses and their respective correlation to clinical outcomes. Plasma from ICU patients with *Streptococcus pyogenes* bacteremia (n=13) were included as a control.

**Table 1.** Patient characteristics of ICU patients with *S. aureus* bacteremia.

| Characteristic | ICU patients (n=36) |
| --- | --- |
| Male, n (%) | 26 (72) |
| Age, years mean (SD) | 61 (16) |
| Admission type, n (%) |  |
| Medical | 21 (58.3) |
| Planned surgical | 6 (16.7) |
| Emergency surgical | 9 (25) |
| Site of infection, n (%) |  |
| Pulmonary | 4 (11.1) |
| Cardiovascular | 9 (25) |
| Musculoskeletal | 7 (19.4) |
| Surgical site | 4 (11.1) |
| Abdominal | 2 (5.6) |
| Soft tissue | 6 (16.7) |
| Unknown | 4 (11.1) |
| Comorbidities prior to ICU admission, n (%) |  |
| Diabetes mellitus | 8 (22.2) |
| Non-metastatic solid tumor | 5 (13.9) |
| APACHE IV score, median (range) | 84 (44-162) |
| Septic shock/sepsis, n (%) | 14 (38.9) |
| Length of stay ICU in days, mean (SD) | 18 (19) |
| ICU mortality, n (%) | 6 (16.7) |

Our analysis focused on WTA-specific IgM and IgG2 responses using a similar setup as in the healthy donor group. We observed that IgG2 responses to any of the three WTA glycotypes were comparable between healthy donors and both patient groups (Figure 3a). In contrast, *S. aureus* bacteremia patients exhibited significantly reduced IgM levels to all WTA glycotypes compared to healthy donors, with undetectable WTA-specific IgM in 7 out-of-36 patients (19%) (Figure 3b). Notably, this reduction WTA-specific IgM responses was not observed in patients with *S. pyogenes* bacteremia (Figure 3b), indicating a pathogen-specific phenomenon. While total IgM titers were generally lower in *S. aureus* bacteremia patients compared to healthy controls, they were similar to *S. pyogenes* bacteremia patients (Supplementary Figure 3a, b) and did not correlate with WTA-specific IgM responses (Supplementary Figure 3c). These findings suggest that the reduced WTA-specific IgM antibodies in *S. aureus* bacteremia patients cannot be attributed to a general decrease in overall IgM titers.

**Figure 3.**
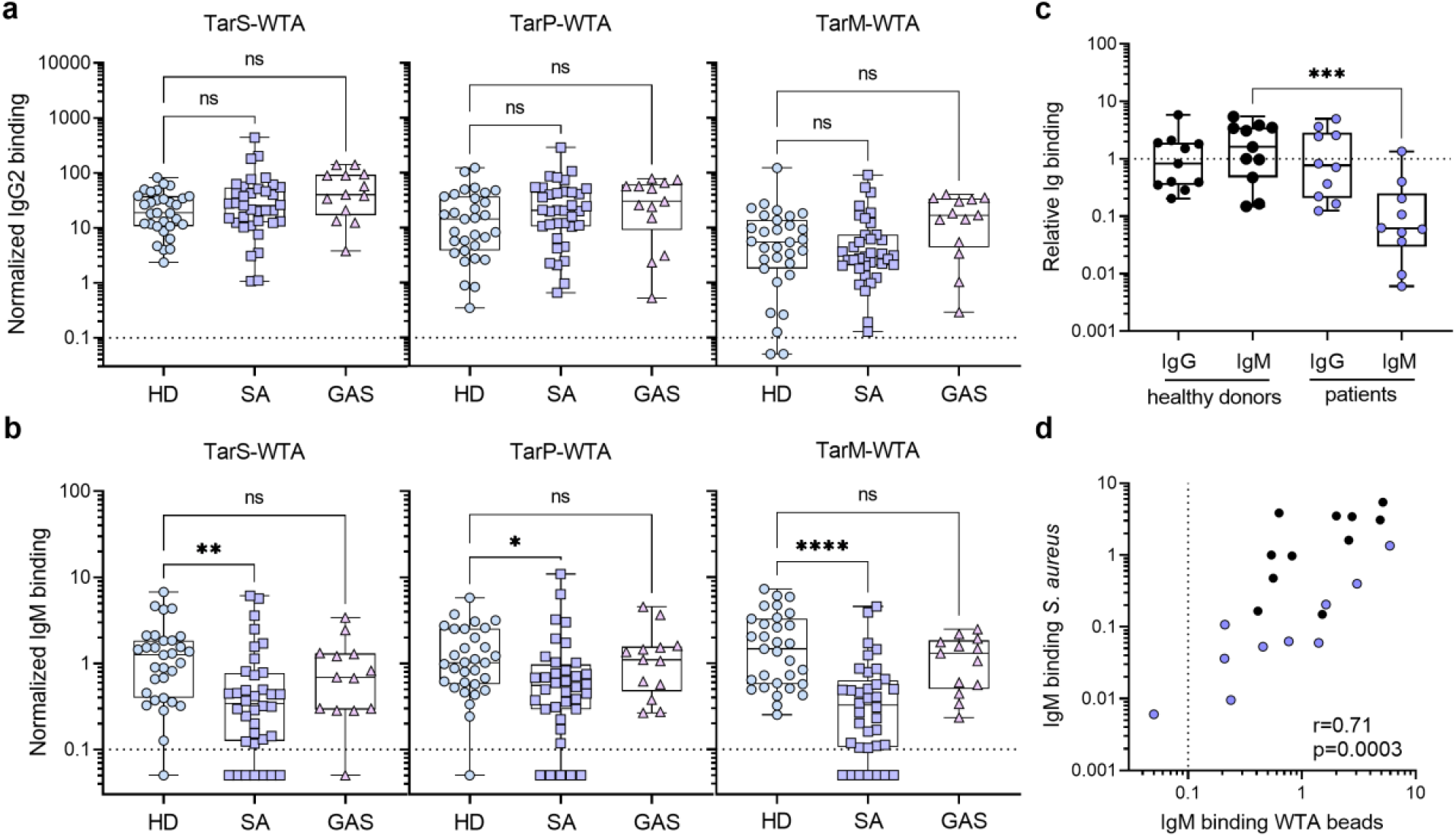
*S. aureus* bacteremia patients display reduced WTA-specific IgM antibody levels. a, b) Normalized binding of (a) IgG2 and (b) IgM to beads coated with TarS-WTA, TarP-WTA and TarM-WTA. Boxplots represent data for healthy donors (HD, n=31) and ICU patients with *S. aureus* (SA, n=36) or *S. pyogenes* (GAS, n=13) bacteremia, and extend from the 25th to 75th percentiles. The line inside the box represents the median, with symbols representing individual donors. c) IgG and IgM binding to *S. aureus* strain Newman Δ*spa*/*sbi* in sera from healthy donors (n=11) and patients with *S. aureus* infection (n=10), defined as relative binding compared to HPS. d) Spearman correlation between IgM binding to Newman Δ*spa*/*sbi* and cumulative IgM binding to TarS- and TarM-WTA beads, patients (n=10) are shown in blue and healthy donors (n=11) in black. Dotted line represents the lower limit of detection, symbols shown below line present extrapolated values. ns = non-significant, *p < 0.05, **p < 0.01, ****p < 0.0001.

WTA is only one of many antigenic molecules on the *S. aureus* surface. To examine the impact of reduced WTA binding on IgM binding to *S. aureus* bacteria as a whole, we utilized sera from a separate cohort of patients with *S. aureus* bacteremia. We compared IgM antibody binding to WTA beads with binding to intact *S. aureus* strain Newman (*tarS/*t*arM* positive; Δ*spa/sbi*). Our analysis revealed significantly lower IgM binding to intact *S. aureus* in sera from patients with *S. aureus* bacteremia compared to healthy donors, while no differences were observed for IgG binding (Figure 3c). Additionally, we found a significant positive correlation between IgM binding to WTA beads and intact *S. aureus*, suggesting that WTA is a major target for IgM on the *S. aureus* surface (Figure 3d).

### WTA-specific IgM levels are inversely associated with disease mortality risk

To examine the association between WTA-specific IgM levels and clinical outcome, specifically mortality, in patients with *S. aureus* bacteremia, we analyzed the antibody responses of the ICU patient cohort. Among the 36 patients in the cohort, 6 (17%) died while in the ICU (Table 1). Stratifying the antibody responses based on mortality revealed that IgM levels, but not IgG2 levels, against TarS- and TarM-modified WTA were significantly lower in deceased patients (Figure 4a, b). For TarP-modified WTA, only IgG2 levels showed a significant reduction in deceased patients (Figure 4a), whereas IgM responses were not significantly different (Figure 4b). There were no significant differences in total IgM or IgG levels, age, length of ICU stay, or underlying co-morbidities between surviving or deceased patients (Supplementary Figure 3d-e, 4).

**Figure 4.**
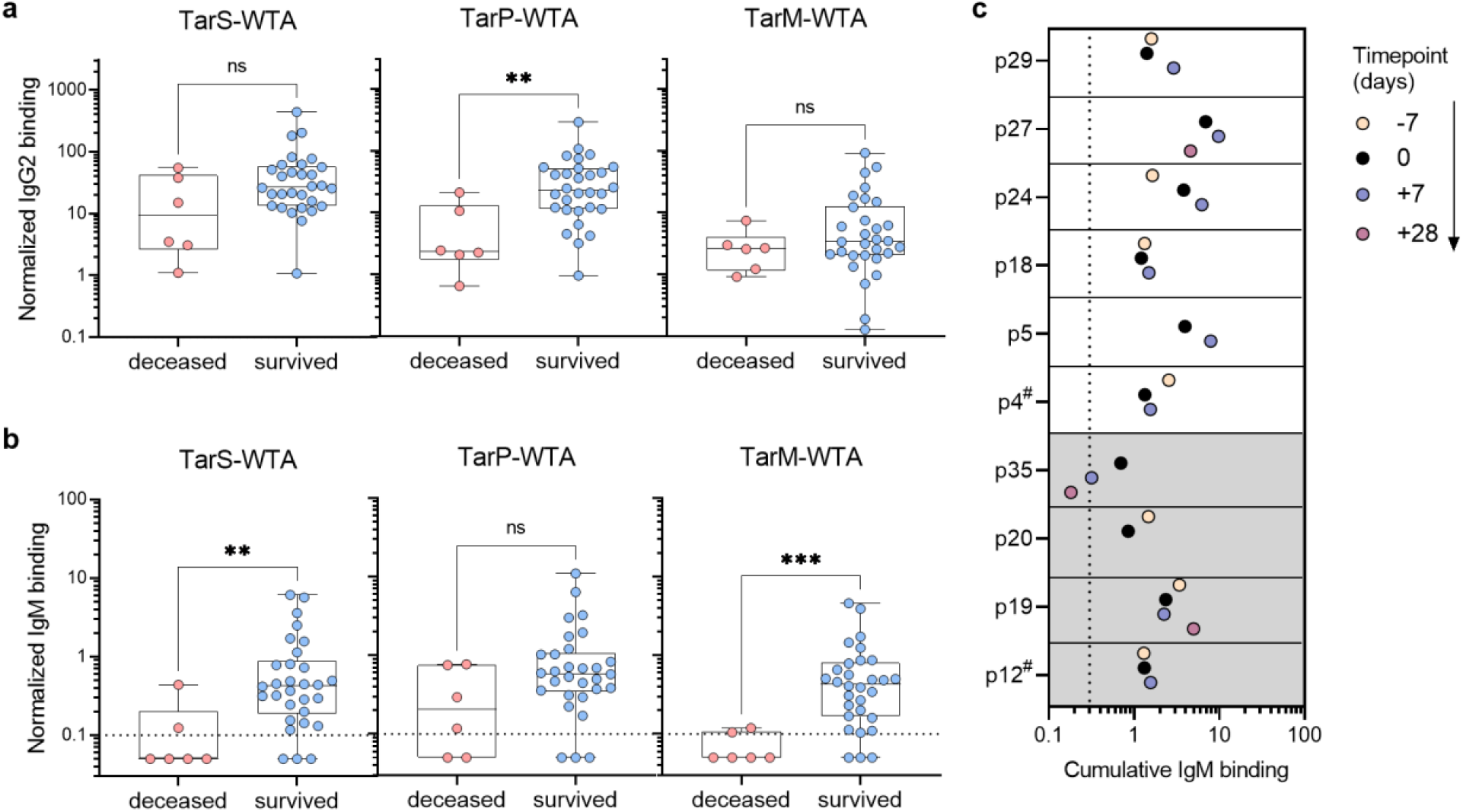
Association between longitudinal WTA-specific antibody responses and ICU mortality in *S. aureus* bacteremia patients. a, b) Normalized binding of plasma-derived WTA-specific a) IgG2 and b) IgM according to the three WTA glycotypes from ICU patients with *S. aureus* bacteremia, stratified by ICU mortality. Symbols indicate individual patients with *S. aureus* infection (n=36). c) Cumulative IgM binding for the three WTA glycotypes (TarS-, TarP- and TarM-WTA) in longitudinal plasma samples (in days from positive blood culture) for ten *S. aureus* bacteremia patients. The shading indicates patients that died on the ICU (p12, p19, p20 and p35). # refers to a different timing of sample acquisition (not -7 days). Dotted line represents the lower limit of quantification, symbols shown below line present extrapolated values. ns= non-significant, *p < 0.05, **p < 0.01.

To determine if the reduced WTA-IgM reactivity observed in *S. aureus* patients was a consequence of systemic infection, we analyzed longitudinal samples from ten patients taken approximately one week before, one week after, and four weeks after the positive blood culture (dependent on sample availability). While a few patients showed minor fluctuations in WTA-specific IgM levels up or down over the course of infection (e.g. patients 24 or 35), the responses to different WTA glycotypes remained relatively constant over time (Figure 4c, Supplementary Figure 5). Notably, for two patients (indicated by #), the available earlier time point was not one week but three years (patient 4) and three months (patient 12) prior to the initial tested sample. These findings suggest that WTA-specific IgM responses are stable over time. Moreover, three patients who later succumbed to *S. aureus* bacteremia already exhibited low WTA-specific IgM responses before the onset of bacteremia, indicating that low WTA-IgM reactivity predisposes to poor disease outcome.

### Low WTA-IgM reactivity impairs complement deposition on live *S. aureus* bacteria

Our findings suggest that low WTA-specific IgM responses are associated with increased disease mortality in patients with *S. aureus* bacteremia. Since IgM-mediated complement activation is not affected by SpA and plays a critical role *S. aureus* killing (figure 2), we hypothesized that impaired bacterial opsonization due to low WTA-specific IgM responses could contribute to this association. To align the patient’s antibody response with the WTA glycoprofile of the infecting *S. aureus* strain, we determined the genetic presence of *tar* glycosyltransferases and the pattern of WTA glycoslyation in available clinical isolates (20). Confirmation of the expressed glycoprofile with GlcNAc-WTA specific Fab fragments is key, since we have recently shown that some *S. aureus* strains harbor inactivating mutations of the encoded Tar enzymes (20). We obtained original isolates from eight patients within the *S. aureus* ICU cohort; four that survived the infection (patient 4, 10, 18 and 27) and four that passed away in the ICU (patient 12, 16, 20 and 35). All eight *S. aureus* isolates contained *tarS* and expressed the corresponding β-1,4-GlcNAc moiety on their surface (Figure 5a); six isolates also expressed *tarM* and corresponding surface expression of α1,4-GlcNAc-WTA (Figure 5a). Of the six patients that had been infected by *tarS*/*tarM*-expressing *S. aureus* strains, four survived the infection (patient 4, 10, 18 and 27) and two passed away (patient 16 and 35) (Figure 5a). IgM binding, C1q- and C3b-deposition on the surface of a *tarS*/*tarM*-positive *S. aureus* strain (Newman Δ*spa/sbi*) were significantly reduced when incubated with plasma from deceased patients compared to surviving patients (Figure 5b, d, e). IgM binding to *S. aureus* strongly correlated with IgM binding to WTA beads (Figure 5c), reinforcing the importance of glycan-specific IgM binding and subsequent complement C3b deposition on the *S. aureus* surface for patient survival following bloodstream infection.

**Figure 5.**
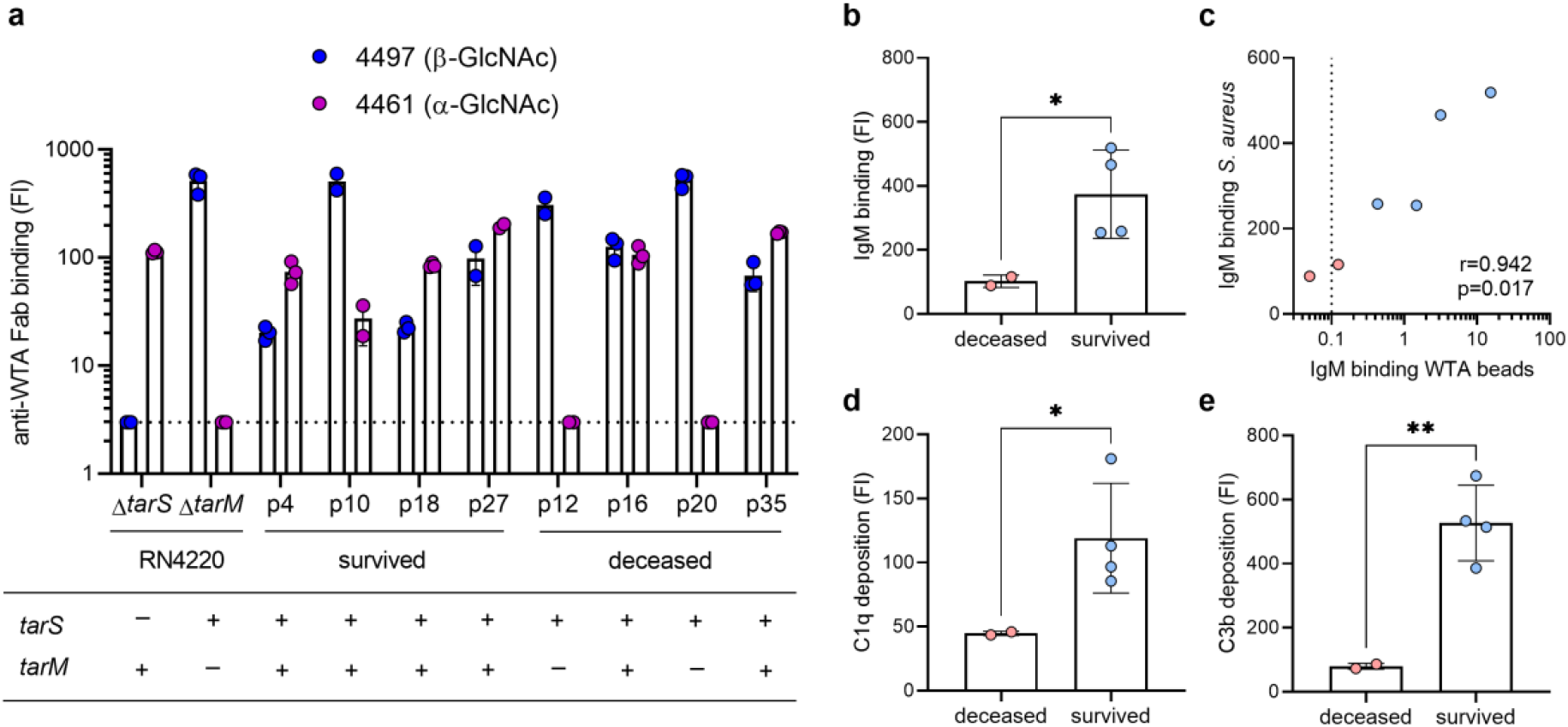
Complement deposition on *S. aureus* by WTA glycoprofile-specific IgM is reduced in ICU patients that succumbed to *S. aureus* bacteremia. a) Expression of β-GlcNAc-WTA (Fab clone 4497) and α-GlcNAc-WTA (Fab clone 4461) by clinical isolates from eight patients (survived patients: p4, p10, p18, p27; deceased patients: p12, p16, p20, p35) and reference strains RN4220 Δ*tarS* and Δ*tarM*. Below the corresponding WTA glycosyltransferase genotypes (*tarS*, *tarM*) as determined by PCR analysis. The dotted line indicates background staining. b) IgM binding to *S. aureus* strain Newman Δ*spa*/*sbi* in 1% plasma from six patients infected with a *tarS*^+^*tarM*^+^ *S. aureus* isolate (as shown in a), stratified on ICU mortality. c) Spearman correlation between IgM binding to *S. aureus* (shown in panel a) and cumulative IgM binding to TarS-WTA and TarM-WTA beads in six patients, with deceased patients shown in red and survived patients in blue. Dotted line represents the lower limit of detection, symbols shown below line present extrapolated values. d, e) Deposition of (d) C1q and (e) C3b on Newman Δ*spa*/*sbi*, pre-opsonized with 3% patient plasma, stratified on ICU mortality. Anti-WTA Fab staining, deposition of IgM, C1q and C3b on *S. aureus* bacteria is depicted as geometric mean fluorescence intensity (FI) (mean + SD of biological duplicates or triplicates), each symbol indicates individual patients. *p < 0.05, **p < 0.01.

## Discussion

In this study, we conducted antibody profiling of glycosylated *S. aureus* RboP-WTA, a prominent glycopolymer expressed by nearly all *S. aureus* strains (20, 28). For this work, we took advantage of synthetic glycosylated RboP-WTA oligomers, which resemble bacteria-expressed WTA as previously confirmed by WTA-specific monoclonal antibodies and recognition by the C-type lectin receptor langerin (15, 29). Our data confirm the near-universal presence of *S. aureus*-specific antibodies, including both IgM and IgG targeting WTA (14, 22), in healthy individuals, confirming widespread exposure to the bacterium in the absence of clinical disease (15). Remarkably, IgM outperformed IgG in complement activation and conferred protection against systemic *S. aureus* infection in mice, while being unaffected by the major *S. aureus* virulence factor SpA. Importantly, WTA-specific IgM levels, but not IgG or IgA, were significantly lower among *S. aureus* bacteremia patients, particularly in non-survivors, and correlated with reduced complement activation, a vital response for recruiting and activating neutrophils (6, 30). Our findings suggest that IgM plays a critical role in protecting against severe *S. aureus* infections, challenging the notion that it is merely a transitional antibody isotype. Further studies in different patient cohorts, for example children or patients with *S. aureus* pneumonia or severe skin and soft tissue infection, are needed to determine the generalizability of our findings and to investigate if insufficient IgM antibodies targeting WTA (or other surface antigens) likewise represents a risk factor for poor clinical outcome.

Our findings may have important implications for clinical care management, particularly in risk assessment and prophylaxis for hospitalized patients. Previous case reports have indicated that patients with primary selective IgM deficiency are more susceptible to *S. aureus* infections (31, 32), and reduced IgM responses can also occur in individuals with advancing age or impaired splenic function (33, 34). Based on our data, low WTA-specific IgM titers are a risk factor for development of *S. aureus* bacteremia and poor clinical outcomes. Conceivably, patients with a planned procedure that holds risk for opportunistic *S. aureus* infections could be evaluated for WTA-specific IgM titers, similar to the assessment of other risk factors such as neutrophil counts and *S. aureus* colonization (35, 36), which clinicians could weigh in decisions on antibiotic selection or frequency of monitoring. Moreover, future studies could explore the potential prophylactic or therapeutic application of IgM to improve outcomes in *S. aureus* bacteremia, for example using IgM-enriched IVIG (37, 38). And while greatly underexplored compared to IgG subtypes, preclinical proof-of-principle studies have been reported for pathogen-targeted therapeutic monoclonal IgM antibodies against influenza (39), SARS-CoV2 (40), and *Neisseria meningitidis* (41).

The potential advantages of opsonic IgM over IgG in protecting against *S. aureus* bacteremia has implications for vaccine development. Current vaccination strategies for *S. aureus* have focused on inducing high levels of IgG, the classic opsonic antibody, but these approaches have consistently failed in human clinical trials, despite promising results in pre-clinical animal studies. The effectiveness of IgG-mediated strategies may be hindered by the presence of SpA, which effectively interferes with IgG-mediated downstream functions (excluding IgG3). Our data from a cohort of *S. aureus* bacteremia patients suggest that efforts to boost durable opsonic IgM responses through vaccination may be more successful, partly because SpA does not block IgM-mediated effector functions like it does with IgG. Characterizing the B cells responsible for producing these glycan-specific IgM antibodies, as well as understanding how to induce their production, will advance our fundamental understanding of protective adaptive immunity in humans. These new insights could likewise inform vaccine development efforts for other human bacterial pathogens that express IgG-Fc or IgA-Fc binding immune evasion factors on their surfaces, including *S. pyogenes* (42).

In conclusion, we provide evidence of a key role for opsonic IgM in protective immunity to *S. aureus*. The manipulation of B cell responses by pathogens to their own advantage is an area of increasing research interest (43), including the observation that Fc-binding by SpA *in vivo* can trigger large-scale supraclonal B-cell depletion by VH-targeted activation-induced cell death in mouse models (44, 45). Further research to dissect the host-pathogen interactions dictating opsonic IgM responses to *S. aureus* can contribute to risk stratification of hospitalized patients and/or rational design of antibody-based therapies and vaccines against this foremost human pathogen.

## Materials and methods

### Sample collection – Ethics statements

Blood from healthy individuals (n=31) was collected in EDTA tubes with full informed consent and approval from the Institutional Review Board of the University Medical Center Utrecht (METC protocol 07-125/C, approved March 1, 2010) in accordance with the Declaration of Helsinki. EDTA-plasma was obtained by centrifugation (10 min, 2,000x*g* at 4°C), and stored at -80°C.

Patients, admitted to the intensive care unit (ICU) were included based on specific inclusion criteria as part of the Molecular Diagnosis and Risk Stratification of Sepsis (MARS) study (ClinicalTrials.gov, NCT01905033). The Institutional Review Board approved an opt-out consent method (protocol number 10-056C). Plasma samples were collected from leftover blood drawn for routine care in EDTA-treated tubes on day one after a positive *S. aureus* (n=36) or *Streptococcus pyogenes* (n=13) blood culture and stored at -80°C within 4 hours after collection.

Blood from patients with *S. aureus* bacteremia (n=10, METC protocol 19/495) and healthy donors (n=11, METC protocol 07-125/C) were collected in serum tubes and centrifuged for 10 min at 3,000 rpm, 4°C. Serum samples were aliquoted and stored at -80°C. Samples were received within one hour of blood draw and processed immediately.

### Glycosylation of synthetic RboP-WTA hexamers and coating to beads

Biotinylated RboP-WTA hexamers were synthesized and glycosylated using recombinant TarS, TarP, and TarM enzymes as previously described (15, 19, 29, 46). Briefly, biotinylated RboP oligomers (0.17 mM) were incubated for 2 h at room temperature with the respective recombinant glycosyltransferase enzymes (6.3 µg/ mL) and UDP-GlcNAc (2 mM, Merck) in glycosylation buffer (15 mM HEPES, 20 mM NaCl, 1 mM EGTA, 0.02% Tween 20, 10 mM MgCl2, 0.1% BSA, pH 7.4) (Supplementary Figure 1a). The glycosylated RboP hexamers were then coupled to magnetic streptavidin (5 x 10^7^) beads for 15 min at room temperature (Dynabeads M280 Streptavidin, Thermo Fisher Scientific). The coated beads were washed three times with PBS 0.1% BSA 0.05% Tween-20 (PBS-BT) and stored at 4°C. The successful coating of WTA beads was validated by binding of monoclonal IgG1 antibodies (3 µg/ml) specific for α-GlcNAc-WTA (TarM-modified WTA, clone 4461), β-GlcNAc WTA (TarS- and TarP-modified WTA, clone 4497) and β-1,4-GlcNAc WTA (TarS-modified WTA, clone 6292), followed by detection with goat-anti human kappa-Alexa Fluor 647 (5 µg/ml, Southern Biotech) and analysis by flow cytometry (BD FACSVerse).

### Production of recombinant monoclonal antibodies

Monoclonal antibodies (mAbs) targeting β-GlcNAc-WTA (clone 4497) in different Ig isotypes and IgG subclasses were expressed in EXPI293F cells (Thermo Fisher) as previously described (29, 47),. Heavy chain (hG) and kappa light chain (hK) constant regions for human IgG1, IgG2, IgG3, IgM and IgA1 were cloned into the XbaI-AgeI cloning sites of the pcDNA34 vector (ThermoFisher), (Supplementary Table 1. along with previously described variable regions (VH and VL) derived from patent WO 2014/ 193722 A1.50 (14, 25). The VH and VL sequences, preceded by a Kozak sequence (ACCACC) and the HAVT20 signal peptide (MACPGFLWALVIST-CLEFSMA), were codon-optimized for human expression and synthesized as gBlocks (IDT). Gibson assembly was used to clone VH and VL gBlocks into the pcDNA34 vector, upstream of the Ig heavy chain (hG) and kappa light chain (hK) constant regions, following the manufacturer’s instructions. NheI and BsiWI were used as the 3′ cloning sites for VH and VL, respectively, to preserve the amino acid sequence of the immunoglobulin heavy and kappa light chains nce. For IgM, BamHI was used as the 3’ cloning site for VH. The constructs were transformed into E. coli TOP10F′ cells through heat shock, and clones were verified by PCR and Sanger sequencing (Macrogen). Plasmids were isolated using the NucleoBond Xtra Midi kit (Macherey-Nagel) and sterilized using 0.22 μm Spin-X centrifuge columns (Corning). For protein production, we used EXPI293F cells and their expression medium (Thermo Fisher); cells were culturedat 37°C, 8% CO2 in conical flasks with culture filter caps (Sigma) placed on a rotation platform (125 rotations/min). One day prior to transfection, the cells were diluted to a concentration of 2 x 10^6^ cells/mL, and 100 mL of cell culture was used for transfection the next day. In 10 mL of Opti-MEM (Thermo Fisher), 500 µL PEI-max (1 µg/µL; Polysciences) was mixed with DNA (1 µg/mL cells) in a 3:2 ratio of hK and hG vectors. After a 20 min incubation at room temperature, this DNA/PEI mixture was added dropwise to 100 mL of EXPI293F cells (2 × 10^6^ cells/mL). After 5 days, Ig expression was verified by SDS-PAGE, and the cell supernatant was collected by centrifugation and filtration through a 0.45 μM filter. IgG1 and IgG2 were purified using a HiTrap Protein A column (GE Healthcare) and Äkta Pure (GE Healthcare). Elution was performed in 0.1M citric acid, pH 3.0, and neutralization done with 1M Tris, pH 9.0. IgG3 was purified using a HiTrap Protein G column (GE Healthcare), with elution in 0.1M Glycine-HCl, pH 2.7, followed by neutralization with 1M Tris, pH 8.0. IgM purification involved dialysis against PBS, and additional NaCl was added to the IgM preparation to achieve a final concentration of 500 mM before application to a POROS™ CaptureSelect™ IgM Affinity matrix (Thermo Scientific) column. IgM was eluted using 0.1M Glycine-HCL pH 3.0 on the ÄKTA Pure system. 0.5M NaCl was added to the pooled fraction, which was then neutralized with 1M Tris pH 7.5. For IgA1 purification, a Jacalin agarose (Thermo scientific) column was used, followed by elution with 0.1M Melibiose (Sigma). All Ig fractions underwent overnight dialysis in PBS at 4°C, and purified mAbs were stored at −20°C.

### Antibody binding to WTA beads

To analyze the presence of WTA-specific antibodies in human plasma or sera, we incubated 1 x 10^5^ beads coated with glycosylated RboP-WTA oligomers (WTA beads) and non-coated beads with a three-fold serial dilution range of human plasma or sera (concentrations ranging between 0.01% - 3%) in a 96-well round bottom plate (Greiner) at 4°C in PBS-BT for 20 minutes. The beads were washed once with PBS-BT using a plate magnet, incubated with a mixture of either (1) goat anti-IgG-PE, goat anti-IgM-FITC and goat F(ab)2 anti-IgA-Alexa Fluor 647 or (2) mouse anti-IgG1 Fc-PE, anti-IgG2 Fc-Alexa Fluor 488 and mouse anti-IgG3 hinge-Alexa Fluor 647 (1 µg/ml, all from Southern biotech) for an additional 20 minutes at 4°C. After another wash, the beads were analyzed by flow cytometry (BD FACSVerse) (Supplementary Figure 1b).

### Data analysis of antibody binding to WTA beads

Antibody binding to WTA beads was assessed by flow cytometry in triplicate or duplicate. The geometric mean fluorescence intensity (geoMFI) values were corrected for median background binding to non-coated beads, except for sera analysis from healthy donors and patients (Figure 2d). Background-corrected geoMFI values were interpolated using a standard curve of β-GlcNAc WTA specific mAb (clone 4497 in IgG1/IgG2/IgG3/IgM/IgA isotype, 0.003 - 10 µg/ml) binding to TarS-WTA beads (Supplementary Figure 1c). For longitudinal sample analysis, ββ-1,4-GlcNAc WTA coated beads (46) were used for the 4497-mAb standard curve. Interpolated values were adjusted for the dilution factor, and the mean normalized antibody binding was calculated using values from at least two dilutions. Values equal to or lower than background were assigned a value of 0.05. Pooled human EDTA-plasma from nine healthy donors was included in each measurement as control sample, ensuring an inter-assay coefficient of variation (CV) <25% (Supplementary Figure 1d).

### Bacterial strains and culture conditions

*S. aureus* clinical isolates, including N315 wildtype (WT), N315 Δ*spa*, Newman WT, Newman Δ*spa/sbi*, RN4220 Δ*tarS* and RN4220 Δ*tarM*, were cultured overnight in 3 mL Todd-Hewitt broth (THB; Oxoid) at 37°C with agitation. For Newman Δ*spa/sbi* mAmetrine (10), bacteria were grown in the presence of 10 µg/ml chloramphenicol. The following day, overnight cultures were subcultured in fresh THB and grown until reaching mid-exponential growth phase, corresponding to an optical density of 0.5-0.6 at 600 nm (OD600). Bacteria were collected by centrifugation (10 min @ 3,000g, 4°C) and resuspended in PBS 0.1% BSA to an OD_600_ of 0.4 for WTA glycoprofiling. For complement deposition and antibody binding experiments, bacteria were resuspended in RPMI supplemented with 0.05% human serum albumin (HSA) or 0.1% BSA (RPMI-A) to an OD600 of 0.4-0.5, and stored at -20°C until further use.

### Complement deposition assays

To assess antibody-mediated complement deposition on intact *S. aureus*, bacteria (∼1 x 10^6^ CFU) were incubated with diluted monoclonal antibodies (IgM/IgG2) or human plasma (1:33 and 1:100) in RPMI-A for 30 mins at 4°C. During this incubation, a titration of wild-type recombinant protein A (SpA-WT, 0.15-100 nM), produced as described in (11), was added simultaneously with 4497-IgM (1 nM) or 4497-IgG2 (10 nM). The bacteria were washed, collected by centrifugation, and incubated with 1% IgG- and IgM-depleted human serum (8) for 30 minutes at 37°C in RPMI-A. Subsequently, the bacteria were washed with RPMI-A and incubated with rabbit F(ab’)_2_ anti-human C3c-FITC (also reactive with C3b), rabbit F(ab’)_2_ anti-human C1q-FITC (both at 5 µg/ml, Dako as described in (9)), or goat F(ab’)_2_ anti-human IgM-PE (5 µg/ml, Southern Biotech) for 30 minutes at 4°C. The bacteria were washed, fixed in 1% paraformaldehyde in RPMI, and analyzed by flow cytometry (BD FACSVerse or Canto) and FlowJo (V10.8.1).

### Neutrophil killing assays

Human blood was collected after informed consent from healthy human volunteers as approved by the University of California San Diego (UCSD) Human Research Protection Program. Human neutrophils were freshly isolated using Polymorphprep (Alere technologies) per manufacturer instructions. Bacteria were opsonized with 10 nM 4497-IgM or 10 nM 4497-IgG2 monoclonal antibodies in the presence of 2% baby rabbit serum (Pel Freeze) in RPMI-A for 30 minutes at 37°C while shaking (650 rpm). Subsequently, freshly isolated human neutrophils were added at a ∼1:10 bacteria to cell ratio in triplicate and incubated for 60 min at 37°C with agitation (200 rpm). To release the internalized bacteria, neutrophils were lysed by incubating 15 min on ice with 0.3% (wt/vol) saponin (Sigma-Aldrich) in sterile water. Samples were serially diluted in PBS and plated on THA plates in duplicate. CFUs were counted after overnight incubation at 37°C, and percentage survival was calculated and normalized over inoculum.

### Passive immunization and systemic *S. aureus* infection *in vivo*

Mouse studies were reviewed and approved by the Institutional Animal Care and Use Committee and conducted in accordance with the regulations of the Animal Care Program at University of California, San Diego. Six weeks old female BALB/c mice were purchased from Charles River Laboratories. All mice were housed in specific-pathogen free facilities and age-matched mice were used for *in vivo* experiments.

*S. aureus* N315 WT was grown overnight from a freshly streaked blood agar plate, diluted 1:200 in THB and grown to an OD600 of 0.7, followed by two washes with PBS. Seven to 10-week old BALB/c female mice were anaesthetized with isofluorane and injected intravenously with monoclonal IgM antibodies (30 µg in 150 µl PBS) by retro-orbital injection. After 3 hours, mice were infected with *S. aureus* N315 (3×10^7^ CFU) by intra-peritoneal (i.p.) injection. Spleen and kidneys were harvested after 24 hours post-infection, homogenized in phosphate-buffered saline (PBS), serially diluted and plated on THB agar plates for CFU enumeration.

### IgG and IgM binding to *S. aureus* Newman Δ*spa*/*sbi*

Newman Δ*spa*/*sbi* mAmetrine bacteria were diluted 50 times from frozen stocks (OD600 of 0.5), and incubated with a serial dilution of heat inactivated (56°C 30 min) human sera in RPMI-A for 30 minutes at 4°C at 600 rpm to allow antibody binding. Heat-inactivated human pooled serum was taken as reference, setting binding of this sample to 1. The bacteria were washed, collected by centrifugation (7 minutes at 3,500 rpm, 4°C) and incubated with goat anti-human IgG-Alexa Fluor 647 and goat anti-human IgM-PE (both Southern Biotech) for 30 minutes at 4°C at 600 rpm. The bacteria were washed and fixed in PBS 1% formaldehyde before analysis by flow cytometry (BD FACSVerse) and FlowJo (V10.8.1).

### WTA glycoprofiling of *S. aureus* isolates

DNA was isolated from *S. aureus* isolates, including RN4220Δ*tarS* and RN4220Δ*tarM* by incubation with lysostaphin and achromopeptidase (both at 100 µg/ml, Sigma) in 0.5 M NaCl, 10 mM Tris-HCl pH 8.0 at 37°C for 30 minutes. Samples were boiled for 5 minutes at 100°C, diluted five-fold in 1 mM EDTA, 10 mM Tris-HCl pH 8.0 and stored at -20°C until further analysis. The presence of *tarS*, *tarP*, and *tarM* was determined by PCR analysis using the following primers: tarP (up) 5′-CTTCACGAAAGAGCACTAGAAG-3′ and tarP (dn) 5′-TTCCCGGCAAGTTGGTG-3′, tarS (up) 5′-GTGAACATATGAGTAGTGCGTA-3′ and tarS (dn) 5′-CATAATGTCCTTCGCCAATCAT-3′ and tarM (up) 5’-GGGATACCCATATATTTCAAGG-3’ and tarM (dn) 5’-CAATTCGCTTCGTTGGTACCATTC-3’.

To analyze the correlation between the *tar* genotype and expressed WTA glycoprofile on the *S. aureus* surface, bacteria were stained with Fab fragments (10 µg/ml) specific for α-GlcNAc WTA (clone 4461), β-GlcNAc WTA (clone 4497) as described previously (1), followed by staining with goat F(ab’)_2_ anti-human kappa-Alexa Fluor 647 (5 μg/mL, Southern Biotech), fixation in PBS 1% formaldehyde and analysis by flow cytometry (BD FACSVerse).

### ELISA to determine total IgG and IgM levels in human plasma

To determine total IgG and IgM levels in human EDTA-plasma samples, Maxisorp plates (Nunc) were coated overnight at 4°C with sheep anti-IgG or sheep anti-IgM (2 µg/ml in PBS, ICN Biomedicals). The next day, the plates were washed three times with PBS 0.05% Tween-20 (PBS-T), blocked for 1 h at 37°C with PBS-T containing 4% BSA (Serva). After three washing cycles with PBS-T, plates were incubated for 1 h at 37°C with a concentration range of plasma samples in duplicate (5-fold serial dilution starting at 1:10,000 for IgG, 1:1,000 for IgM) as well as a standard for either IgG (0.56-200 ng/ml, ChromPure human IgG, Jackson Immunoresearch) or IgM (3.12-400 ng/ml, ChromPure human IgM, Jackson Immunoresearch). Following three washing steps, horseradish peroxidase (HRP)-conjugated goat anti-human IgG or goat anti-human IgM (1:6,000, Southern Biotech) was added for 1 h at 37°C, and after washing the plates were developed using tetramethylbenzidine (TMB). After 5-10 minutes, the reaction was stopped by adding 1N H_2_SO_4_, absorbance was measured at 450 nm in an iMark Microplate Absorbance Reader (Bio-Rad), and values were corrected for background signals at 595 nm. Pooled human plasma (n= 9 healthy donors) was included in every measurement as control sample to determine the inter-assay variation, resulting in a coefficient of variation (CV) <25% (data not shown).

### Statistical analysis

Data obtained by flow cytometry was analyzed using FlowJo 10 (FlowJo LLC). Statistical analysis was performed with Prism software (version 8.3; GraphPad). Data between three groups were analyzed using the Kruskal-Wallis test, between two groups the Mann-Whitney U test or unpaired t-test with Welch correction (or Holm-Šídák method for multiple unpaired t-test) was used. Correlations between antibody binding to different WTA beads or *S. aureus* bacteria were assessed using the Spearman correlation. Two-sided p values < 0.05 were considered significant, and are depicted in the figures.

## Acknowledgements

We kindly thank Thilo Stehle for providing recombinant TarP enzyme, Andreas Peschel for the bacterial strains, Alex Stream and Elisabet Bjanes for human neutrophil isolation, Chih-Ming Tsai and Irshad Hajam for support with the in vivo experiments, András Spaan and Pieter-Jan Haas for assistance in obtaining the clinical isolates. We also thank Kok van Kessel, Bart Bardoel, Yvonne Pannekoek, Robin Temming and Rob van Dalen for the valuable discussions.

## Funding and additional information

This work was supported by the Vidi (91713303) and Vici (09150181910001) research program to N.M.v.S. and A.H., which is financed by the Dutch Health Council (NWO), and the BactiVac Training grant for sponsoring A.H. to visit UCSD. The work of P.F.K. was funded by Genmab B.V.

## Notes

### Competing Interest Statement

The authors have declared no competing interest.

### Summary of Updates

ORCID ID link of first author Astrid Hendriks was corrected

