## Supplementary information for "Glycan-specific IgM is critical for human immunity to *Staphylococcus aureus*"

**Supplementary Material**

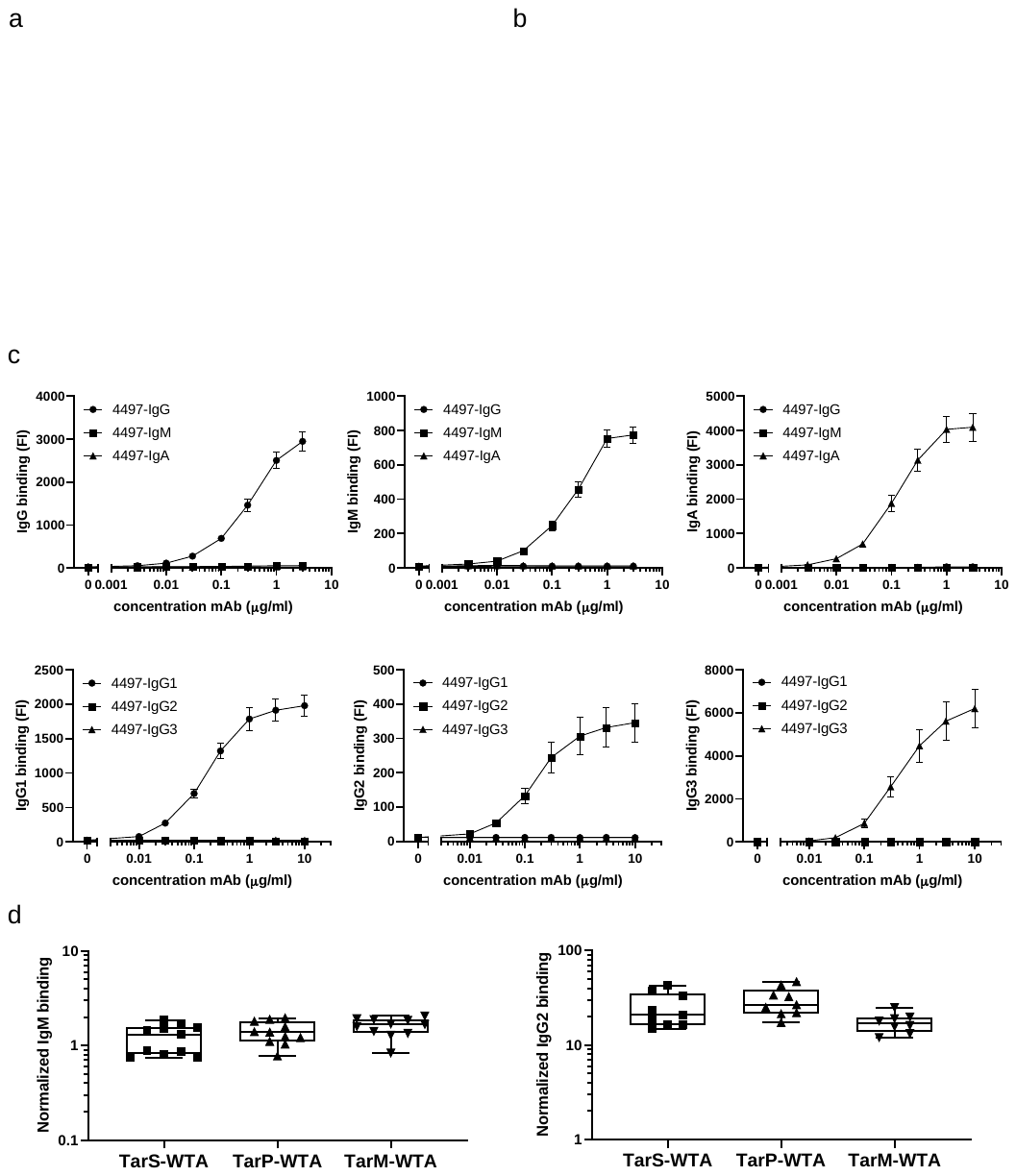

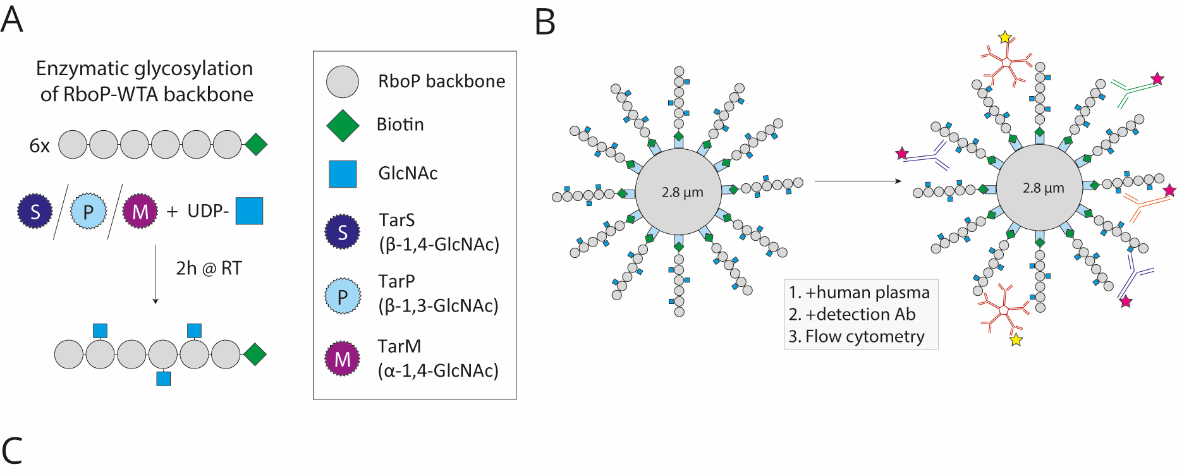

**Supplementary Figure 1** **Schematic overview of semi-synthetic WTA fragments, multiplex set up of bead-based assay and interassay variation.** A) Schematic representation of in vitro enzymatic glycosylation of biotinylated RboP-WTA hexamers by recombinant enzymes (TarS, TarP and TarM), which are then coated to beads (WTA beads). B) General work-flow to measure antibody binding to WTA beads by flow cytometry. WTA beads are incubated with human plasma (step 1), subsequently stained with a combination of three detection antibodies to analyze multiple antibody isotypes/subclasses in a single sample (Step 2) to measure antibody binding by flow cytometry (step 3). C) Binding of β-GlcNAc WTA specific mAb (clone 4497 in different Ig isotypes/IgG subclasses) to TarS-WTA coated beads. Concentration curves were used for data interpolation, data is depicted as geometric mean fluorescence intensity (FI) + standard error for the mean (SEM) of 6 (for IgG subclasses) or 11 (for IgM, IgA and IgG) independent experiments. D) Interassay variation within control sample (pooled plasma, n=9 healthy donors) in normalized IgM and IgG2 binding to TarS-WTA, TarP-WTA and TarM-WTA beads.

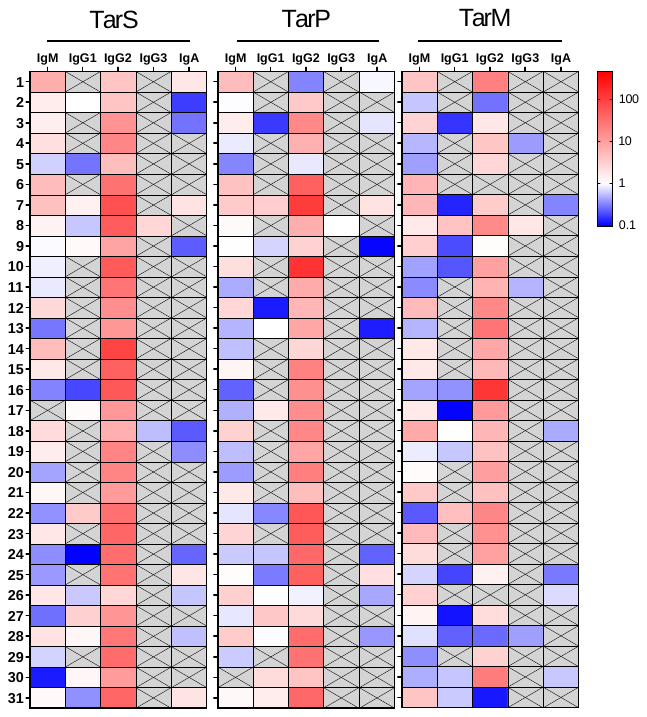

**Supplementary Figure 2 Heatmap of WTA-specific antibody responses in healthy individuals.** Colors match log-transformed normalized binding data for IgM, IgG1, IgG2, IgG3 and IgA to the three WTA glycotypes (mediated by TarS, TarP and TarM). Grey boxes with crosses indicate values equal to or below detection limit (0.1).

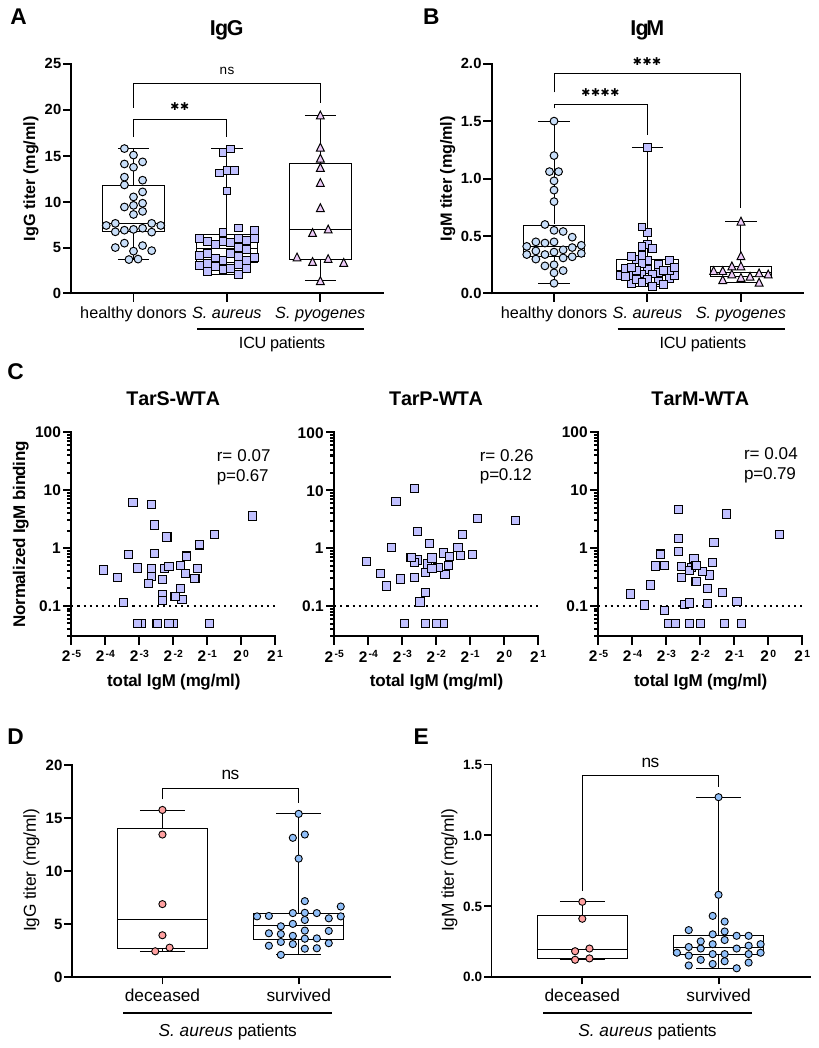

**Supplementary Figure 3 Total IgG and IgM titers in plasma from all included donors and correlation between IgM titer and WTA-specific IgM.** A) IgG titers and B) IgM titers of healthy donors (n=31) and patients with *S. aureus* (n=36) or *S. pyogenes* (n=13) bacteremia. C) Spearman correlation between WTA-specific IgM responses and total IgM titers in patients with *S. aureus* bacteremia, dotted line represents detection limit. D) IgG titers and E) IgM titers in patients with *S. aureus* bacteremia, grouped by ICU mortality. Symbols indicate individual donors, boxplots extend from the 25th to 75th percentiles and the line inside the box represents the median. ***p < 0.001, ****p < 0.0001, ns=not significant.

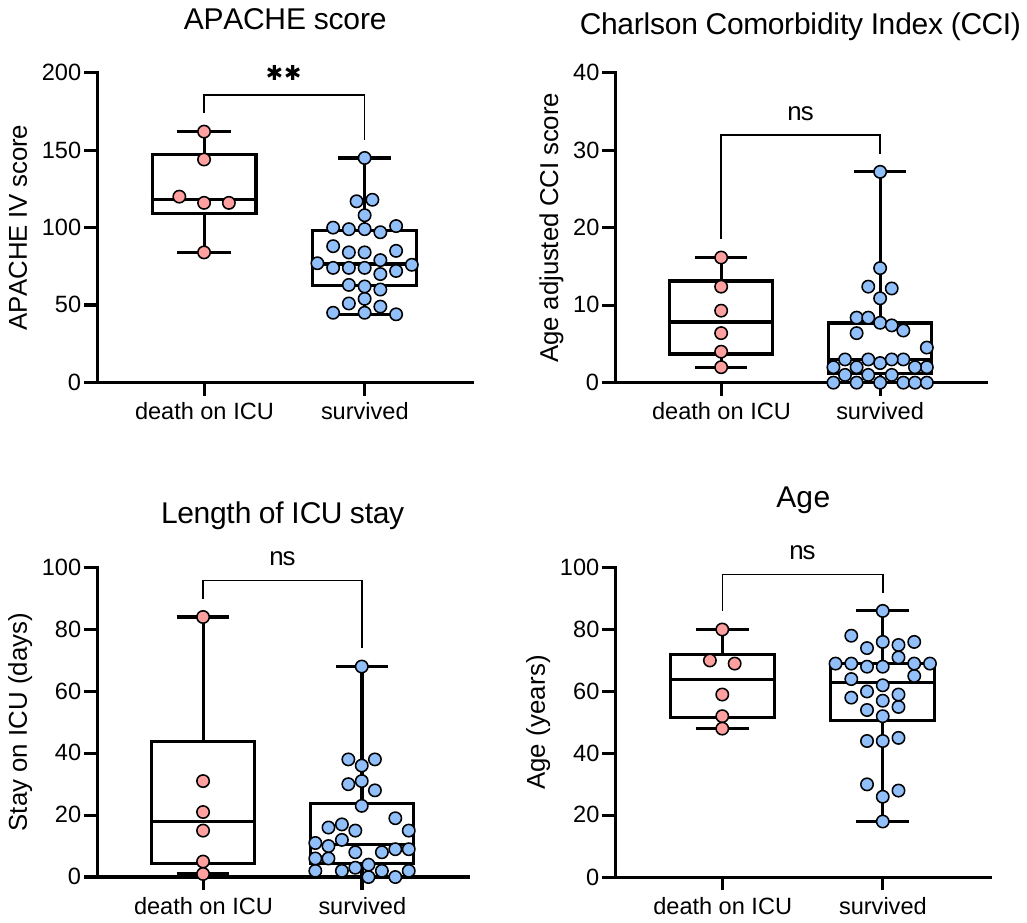

**Supplementary Figure 4 Clinical data ICU patients with *S. aureus* bacteremia**

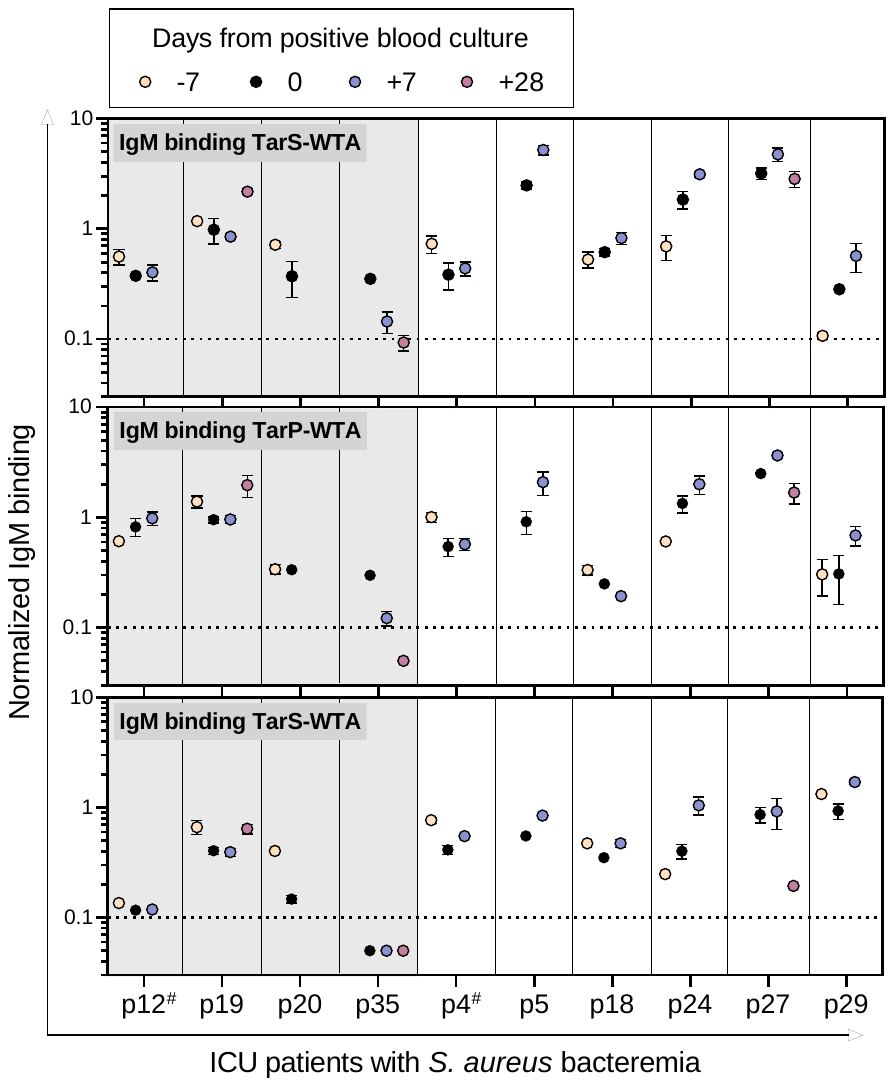

**Supplementary Figure 5 Longitudinal analysis of WTA-specific IgM responses in *S. aureus* bacteremia patients.** Normalized IgM binding for A) TarS-WTA, B) TarP-WTA and C) TarM-WTA at different time points (in days from positive blood culture). Dotted line represents the lower limit of detection, symbols shown below line present extrapolated values.

**Supplementary table 1 protein sequence for 4497 antibody isotypes.**

|  | **Sequence** |
| --- | --- |
| **Variable heavy chain** | |
| Anti-WTA-IgG (4497) | EVQLVESGGGLVQPGGSLRLSCSASGFSFNSFWMHWVRQVPGKGLVWISFTNNEGTTTAYADSVRGRFIISRDNAKNTLYLEMNNLRGEDTAVYYCARGDGGLDDWGQGTLVTVSS |
| **Variable light chain** | |
| Anti-WTA-IgG (4497) | DIQLTQSPDSLAVSLGERATINCKSSQSIFRTSRNKNLLNWYQQRPGQPPRLLIHWASTRKSGVPDRFSGSGFGTDFTLTITSLQAEDVAIYYCQQYFSPPYTFGQGTKLEIK |
| **Constant heavy chain** | |
| IgG1 | ASTKGPSVFPLAPSSKSTSGGTAALGCLVKDYFPEPVTVSWNSGALTSGVHTFPAVLQSSGLYSLSSVVTVPSSSLGTQTYICNVNHKPSNTKVDKKVEPKSCDKTHTCPPCPAPELLGGPSVFLFPPKPKDTLMISRTPEVTCVVVDVSHEDPEVKFNWYVDGVEVHNAKTKPREEQYNSTYRVVSVLTVLHQDWLNGKEYKCKVSNKALPAPIEKTISKAKGQPREPQVYTLPPSREEMTKNQVSLTCLVKGFYPSDIAVEWESNGQPENNYKTTPPVLDSDGSFFLYSKLTVDKSRWQQGNVFSCSVMHEALHNHYTQKSLSLSPGK |
| IgG2 | ASTKGPSVFPLAPCSRSTSESTAALGCLVKDYFPEPVTVSWNSGALTSGVHTFPAVLQSSGLYSLSSVVTVPSSNFGTQTYTCNVDHKPSNTKVDKTVERKCCVECPPCPAPPVAGPSVFLFPPKPKDTLMISRTPEVTCVVVDVSHEDPEVQFNWYVDGVEVHNAKTKPREEQFNSTFRVVSVLTVVHQDWLNGKEYKCKVSNKGLPAPIEKTISKTKGQPREPQVYTLPPSREEMTKNQVSLTCLVKGFYPSDIAVEWESNGQPENNYKTTPPMLDSDGSFFLYSKLTVDKSRWQQGNVFSCSVMHEALHNHYTQKSLSLSPGK |
| IgG3 | ASTKGPSVFPLAPCSRSTSGGTAALGCLVKDYFPEPVTVSWNSGALTSGVHTFPAVLQSSGLYSLSSVVTVPSSSLGTQTYTCNVNHKPSNTKVDKRVELKTPLGDTTHTCPRCPEPKSCDTPPPCPRCPEPKSCDTPPPCPRCPEPKSCDTPPPCPRCPAPELLGGPSVFLFPPKPKDTLMISRTPEVTCVVVDVSHEDPEVQFKWYVDGVEVHNAKTKPREEQYNSTFRVVSVLTVLHQDWLNGKEYKCKVSNKALPAPIEKTISKTKGQPREPQVYTLPPSREEMTKNQVSLTCLVKGFYPSDIAVEWESSGQPENNYNTTPPMLDSDGSFFLYSKLTVDKSRWQQGNIFSCSVMHEALHNRFTQKSLSLSPGK |
| IgM | GSASAPTLFPLVSCENSPSDTSSVAVGCLAQDFLPDSITFSWKYKNNSDISSTRGFPSVLRGGKYAATSQVLLPSKDVMQGTDEHVVCKVQHPNGNKEKNVPLPVIAELPPKVSVFVPPRDGFFGNPRKSKLICQATGFSPRQIQVSWLREGKQVGSGVTTDQVQAEAKESGPTTYKVTSTLTIKESDWLSQSMFTCRVDHRGLTFQQNASSMCVPDQDTAIRVFAIPPSFASIFLTKSTKLTCLVTDLTTYDSVTISWTRQNGEAVKTHTNISESHPNATFSAVGEASICEDDWNSGERFTCTVTHTDLPSPLKQTISRPKGVALHRPDVYLLPPAREQLNLRESATITCLVTGFSPADVFVQWMQRGQPLSPEKYVTSAPMPEPQAPGRYFAHSILTVSEEEWNTGETYTCVVAHEALPNRVTERTVDKSTGKPTLYNVSLVMSDTAGTCY |
| IgA1 | ASPTSPKVFPLSLCSTQPDGNVVIACLVQGFFPQEPLSVTWSESGQGVTARNFPPSQDASGDLYTTSSQLTLPATQCLAGKSVTCHVKHYTNPSQDVTVPCPVPSTPPTPSPSTPPTPSPSCCHPRLSLHRPALEDLLLGSEANLTCTLTGLRDASGVTFTWTPSSGKSAVQGPPERDLCGCYSVSSVLPGCAEPWNHGKTFTCTAAYPESKTPLTATLSKSGNTFRPEVHLLPPPSEELALNELVTLTCLARGFSPKDVLVRWLQGSQELPREKYLTWASRQEPSQGTTTFAVTSILRVAAEDWKKGDTFSCMVGHEALPLAFTQKTIDRLAGKPTHVNVSVVMAEVDGTCY |
| **Constant light chain (kappa)** | |
| IgG1,2,3, IgM, IgA1 | RTVAAPSVFIFPPSDEQLKSGTASVVCLLNNFYPREAKVQWKVDNALQSGNSQESVTEQDSKDSTYSLSSTLTLSKADYEKHKVYACEVTHQGLSSPVTKSFNRGEC |
